# Volatile allosteric antagonists of mosquito odorant receptors inhibit normal odor-dependent behaviors

**DOI:** 10.1101/2020.04.26.062919

**Authors:** Georgia Kythreoti, Nadia Sdralia, Panagiota Tsitoura, Dimitrios P. Papachristos, Antonios Michaelakis, Vasileios Karras, David M. Ruel, Esther Yakir, Jonathan D. Bohbot, Stefan Schulz, Kostas Iatrou

**Affiliations:** Institute of Biosciences & Applications, National Centre for Scientific Research “Demokritos”, 15341 Aghia Paraskevi, Greece; Entomology & Agricultural Zoology, Benaki Phytopathological Institute, 14561 Kifissia, Greece; Department of Entomology, The Robert H. Smith Faculty of Agriculture, Food and Environment, The Hebrew University of Jerusalem, Rehovot 76100, Israel; Institute of Organic Chemistry, Technische Universität Braunschweig, 38106 Braunschweig, Germany

**Keywords:** allosteric regulation, cell surface receptor, 7-helix ligand-gated channel, ion channel, ligand-binding protein, mosquito odorant receptors, ORco co-receptor, volatile organic compounds, evolutionary conservation, mosquito repellents, ligand binding sites

## Abstract

Odorant-dependent behaviors in insects are triggered by the binding of odorant ligands to the variable subunits of heteromeric olfactory receptors. Previous studies have shown, however, that specific odor binding to ORco, the common subunit of odorant receptor heteromers, may alter allosterically olfactory receptor function and affect profoundly subsequent behavioral responses. Here we report on the identification of several antagonists of the odorant receptor co-receptor of the African malaria vector *Anopheles gambiae*, AgamORco, in a small collection of natural volatile organic compounds (VOCs) using a relevant insect cell-based screening platform. Because some of the identified antagonists were previously shown to strongly repel *Anopheles* and *Culex* mosquitoes, here we examined the bioactivities of the identified antagonists against *Aedes*, the third major genus of the Culicidae family. The tested antagonists were found to inhibit the function of *Ae. aegypti* ORco *ex vivo* and repel Asian tiger, *Ae. albopictus*, adult mosquitoes. Specific antagonist binary mixtures elicited higher repellency than single antagonists. Binding competition assays suggested antagonist binding to distinct ORco sites as a likely cause for the enhanced repellence of the blends. These findings demonstrate that a simple screening assay may be used for the identification of allosteric modifiers of olfactory-driven behaviors capable of providing enhanced indoor and outdoor protection against multiple mosquito borne infectious diseases.

## INTRODUCTION

Insect odorant receptors (ORs) are heteromeric ligand-gated ion channels expressed by olfactory receptor neurons (ORNs) inside olfactory sensilla 1-3). Together with odorant binding proteins (OBPs) (4) and odorant degrading enzymes (ODEs) (5) that are produced by accessory cells in the olfactory sensilla (6-8) and secreted in the lymph surrounding the ORNs, ORs constitute the molecular gateway to the olfactory pathway and associated behaviors that are important for survival and reproduction (9). The odorant receptor heteromeric complexes consist of an obligatory and highly conserved subunit, ORco (10-14) and one of many variable ligand-binding ORx subunits (12,15) in as yet undetermined molar ratios. Odorant ligands act either as receptor agonists or antagonists in an ORx-specific manner (16-22). In cell cultures, homomeric ORco channels are formed that are activated by specific ORco agonists (OAs) such as VUAA1 and OrcoRAM2 (23-25).

In previous studies, we have used multiple OBPs of the African malaria mosquito vector *Anopheles gambiae* (AgamOBPs; 26-31) as screening tools for the discovery of natural volatile organic compounds (VOCs) capable of modifying olfactory-mediated behaviors (32,33). This effort resulted in the identification of natural compounds with strong repellent activities against both *Anopheles* and *Culex* mosquitoes (33) suggesting the existence of phylogenetically conserved molecular mechanisms and behavioral outputs in mosquitoes. Further studies revealed that the most potent of the identified repellents acted as allosteric inhibitors of multiple AgamORs in the heteromeric complexes and blocked odorant-specific responses by interacting directly with AgamORco (34). Additionally, we have shown that *An. gambiae* ORx/Orco functional responses elicited by ORx-specific odor agonists were enhanced both in terms of potency and efficacy by one to two orders of magnitude in the presence of an OA (35). These findings also pointed to the induction of conformational rearrangements in ORx ligand-bound ORx/ORco receptor complexes caused by the binding of the OA and resulting in enhanced inward currents into the receptor-expressing cells.

In view of these results and given the previously demonstrated importance of ORco for the functionality of OR heteromers and OR-dependent behaviors (36-43), we have employed the lepidopteran insect cell-based assay toward the rapid detection of potential agonists and antagonists of AgamORco. This system relies on the stable expression of homomeric AgamORco in cells constitutively expressing a luminescence emitting calcium biosensor reporter protein. Here, we are reporting on the screening of a small collection of VOCs of plant, arthropod and bacterial origin for the identification of modulators of AgamORco function. The screening resulted in the identification of several AgamORco-specific antagonists. Considering the high degree of phylogenetic conservation of ORco and its functional relevance, which was demonstrated by our previous findings that natural compounds inhibiting AgamORco activity were capable of repelling at least two mosquito genera, *Anopheles* and *Culex*, we examined whether the identified ORco antagonists were also active against the third major genus of Culicidae mosquitoes, *Aedes*. Two tested antagonists elicited significant inhibition of inward currents mediated by VUAA1 in *Xenopus laevis* oocytes expressing *Ae. aegypti* ORco (AaegORco). Examination of the bioactivity of the identified antagonists and binary mixtures thereof against available laboratory populations of *Ae. albopictus* elicited an avoidance behavior, with some of the mixtures causing anosmia-like effects similar to equivalent doses of DEET. Antagonist binding competition assays against an OA point to the simultaneous binding of antagonists to the OA binding site on ORco and to one or more alternative binding sites as a plausible cause for the observed enhanced activities of the binary mixtures.

## RESULTS

The screening platform employed in this study exploits the property of AgamORco homomers to form functional ligand-gated cation channels in cultured lepidopteran cells (34,35). The constitutively expressed reporter photoprotein Photina™ detects the entry of Ca^2+^ ions into the cell upon AgamORco channel activation. The screening protocol, performed in a 96-well format, involved the sequential addition of a tested compound and the Orco Receptor Activator Molecule, ORcoRAM2, as OA, both at concentrations of 100 μM, to the transformed cells (**Figure 1**).

**Figure 1:**
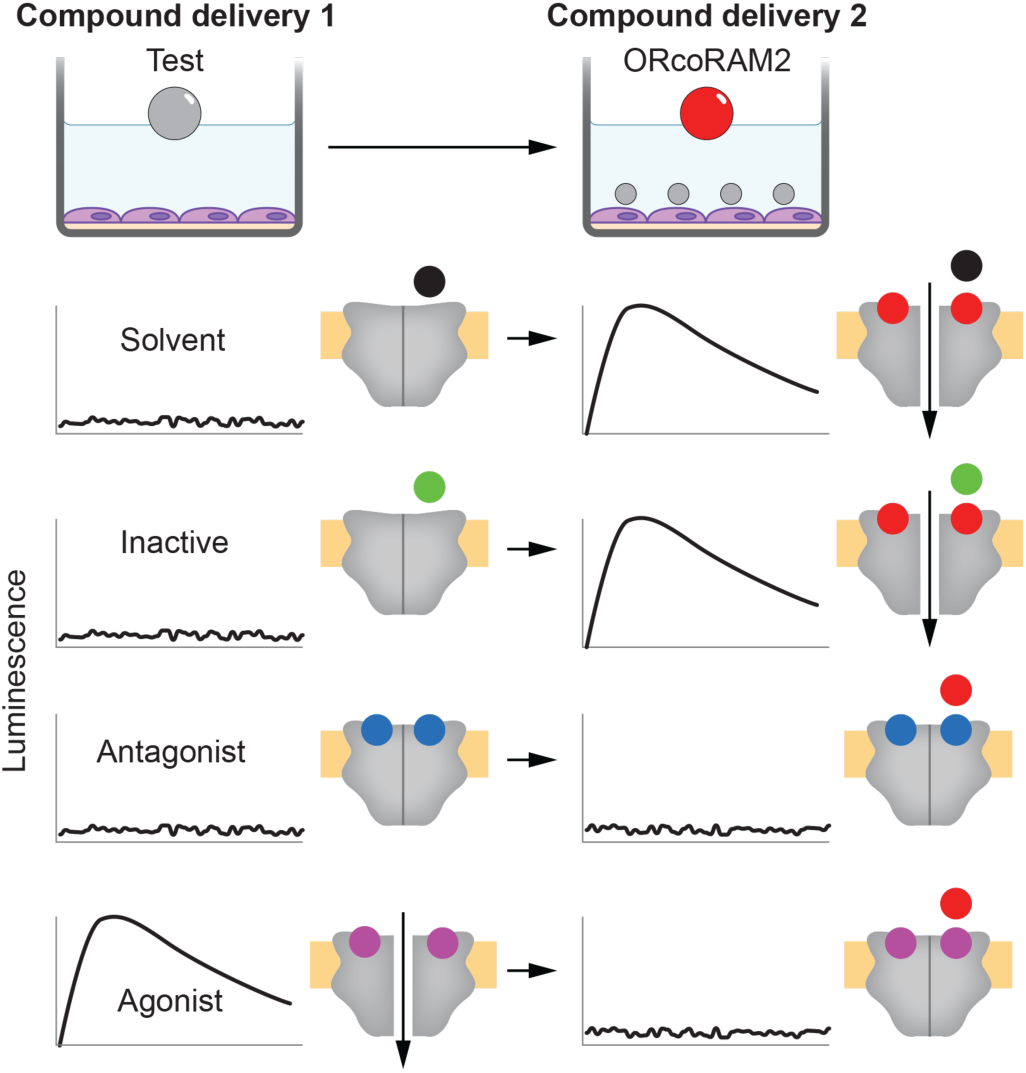
Insect cell-based screening assay for ligand identification. Schematic representation of a two-step screening assay for VOC activity determination performed in a 96-well format containing lepidopteran insect cells expressing *An. gambiae* ORco functional homomeric channel and Photina™ Ca^2+^ biosensor. Initially, a tested VOC is added at a concentration of 100 μM and the response of the ORco channel is monitored. This is followed by addition of 100 μM ORcoRAM2, a known OA, and measurement of the secondary response. The anticipated outcomes and corresponding VOC classifications are indicated. For simplification reasons, the recently deduced homotetrameric structure of the ORco channel is illustrated here as a homodimer. Note also that while the orthosteric binding of antagonists and new agonists in the postulated OA (VUAA1 or OrcoRAM2) site is shown in the Figure, their binding in alternative, allosteric binding sites is also possible but not illustrated here.

Similarly to the working scheme presented earlier (34,35), the general principle for agonist/antagonist screening has been that the addition of the ethanol solvent or a compound devoid of ORco-binding activity would allow the appearance of cellular luminescence upon sequential addition of the ORco agonist, while the addition of a compound acting as an ORco antagonist would prevent, partially or completely, the appearance of luminescence in the cells upon secondary addition of the ORco agonist. The same expression platform also allows the identification of compounds acting as ORco agonists. In that case, addition of an active compound would cause the appearance of cell luminescence, while no response would be expected upon addition of the known agonist after the dissipation of the first luminescence burst due to temporary inactivation of the ORco channel (34).

### Natural VOCs inhibit AgamORco homomeric channel activity

The examination of 50 natural VOCs (**Table S-1**) for the presence of modulators of AgamORco function, employed as control the mosquito repellent IPC (compound II; 44) that was previously shown to be an AgamORco channel antagonist (34). The initial screen resulted in the identification of five hits with AgamORco antagonist activity (**Figure 2**).

**Figure 2:**
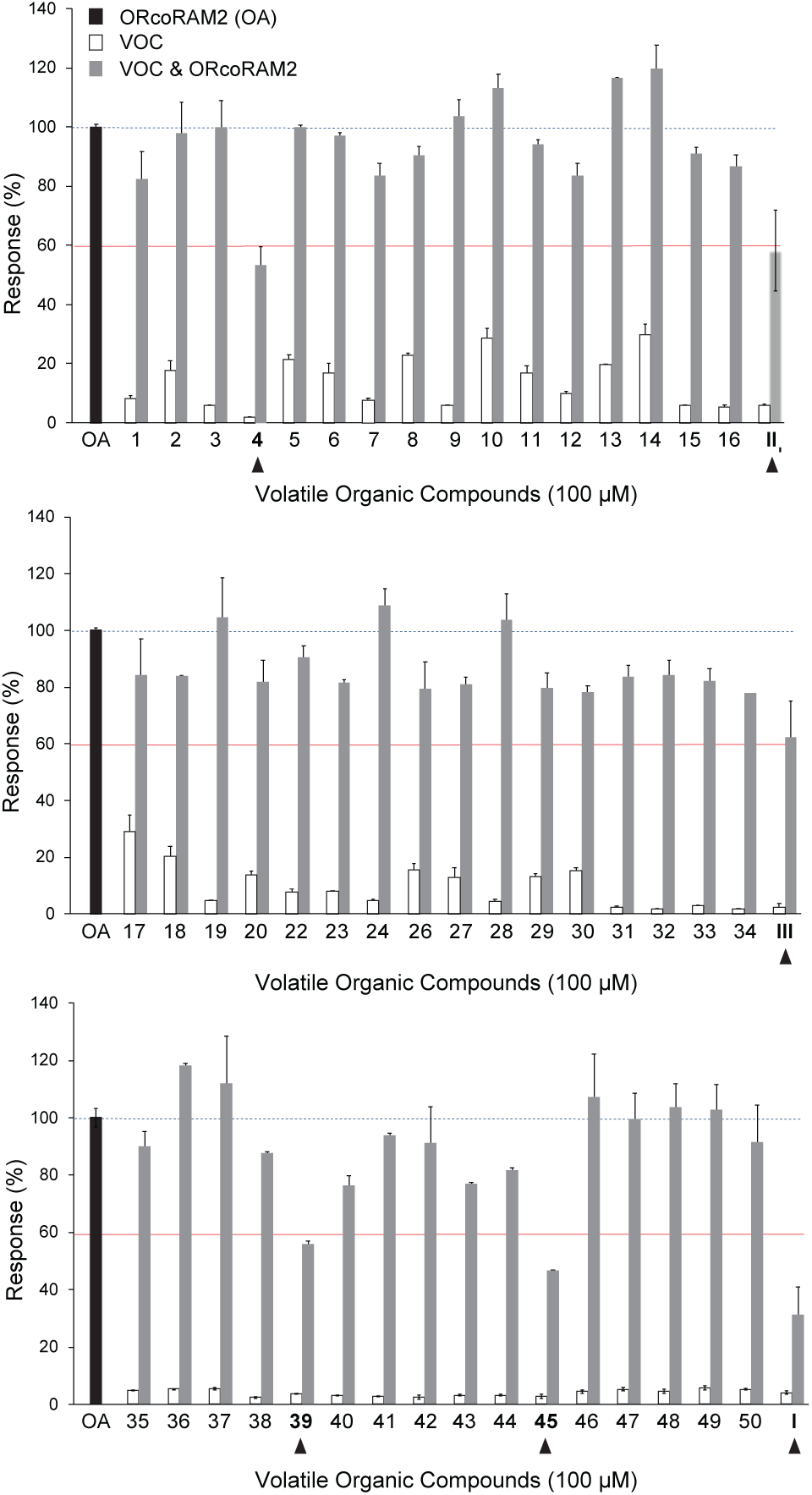
Initial screening results. All compounds were tested at a final concentration of 100μM. The primary compound additions (white bars) do not induce any significant ORco channel function, while secondary additions of the OA (ORcoRAM2) to wells containing previously added, functionally inactive compounds produce responses (grey bars) equal to at least 80% of the full response obtained in the control wells (OA only added, shown with a white bar at right of each panel). ORco antagonist hits produce significantly lower secondary responses, arbitrarily set at ≤60% of the normal channel response, upon OA addition (black bars). Numbers correspond to those of the compounds shown in **Supplemental Table 1**. Error bars indicate mean±SE.

Two of these hits, CRV and CA (compounds I and III, respectively), were previously shown to be effective repellents for *An. gambiae* and *Culex spp* mosquitoes (33) and also to inhibit AgamORco activity to an extent that was not determined at the time (34). In contrast, no relevant information existed concerning the bioactivity of the other three antagonist hits, compounds #4 (LA), #39 (OCT) and #45 (CAR). No AgamORco agonists were observed in this VOC collection.

**Table 1:**
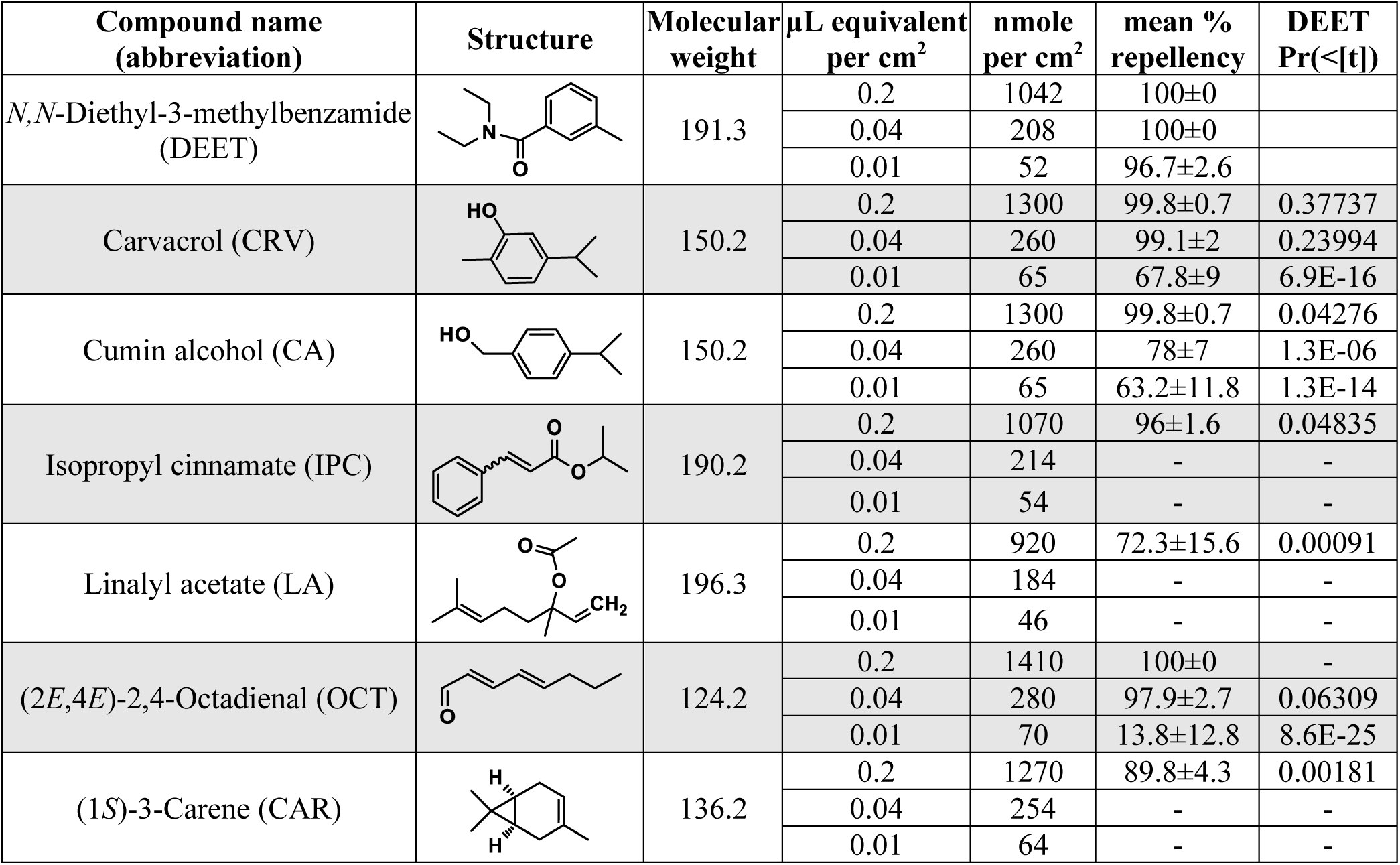
Repellence indices of tested compounds in human hand landing assays. All tests were carried out over a period of 5 minutes. Dichloromethane (DCM) was used as a control and the number of landings for the experiments involving compound equivalent doses of 0.2, 0.04 and 0.01 μL/cm^2^ are presented in **Supplementary Table 2**. Statistically significant differences are those with Pr(<[0.05]). -, not examined.

| Compound name<br>(abbreviation) | Structure | Molecular<br>weight | $\mu\text{L}$ equivalent<br>per $\text{cm}^2$ | nmole<br>per $\text{cm}^2$ | mean %<br>repellency | DEET<br>Pr(<[t]) |
| --- | --- | --- | --- | --- | --- | --- |
| *N,N*-Diethyl-3-methylbenzamide<br>(DEET) |  | 191.3 | 0.2 | 1042 | 100±0 |  |
|  |  |  | 0.04 | 208 | 100±0 |  |
|  |  |  | 0.01 | 52 | 96.7±2.6 |  |
| Carvacrol (CRV) |  | 150.2 | 0.2 | 1300 | 99.8±0.7 | 0.37737 |
|  |  |  | 0.04 | 260 | 99.1±2 | 0.23994 |
|  |  |  | 0.01 | 65 | 67.8±9 | 6.9E-16 |
| Cumin alcohol (CA) |  | 150.2 | 0.2 | 1300 | 99.8±0.7 | 0.04276 |
|  |  |  | 0.04 | 260 | 78±7 | 1.3E-06 |
|  |  |  | 0.01 | 65 | 63.2±11.8 | 1.3E-14 |
| Isopropyl cinnamate (IPC) |  | 190.2 | 0.2 | 1070 | 96±1.6 | 0.04835 |
|  |  |  | 0.04 | 214 | - | - |
|  |  |  | 0.01 | 54 | - | - |
| Linalyl acetate (LA) |  | 196.3 | 0.2 | 920 | 72.3±15.6 | 0.00091 |
|  |  |  | 0.04 | 184 | - | - |
|  |  |  | 0.01 | 46 | - | - |
| (2*E*,4*E*)-2,4-Octadienal (OCT) |  | 124.2 | 0.2 | 1410 | 100±0 | - |
|  |  |  | 0.04 | 280 | 97.9±2.7 | 0.06309 |
|  |  |  | 0.01 | 70 | 13.8±12.8 | 8.6E-25 |
| (1*S*)-3-Carene (CAR) |  | 136.2 | 0.2 | 1270 | 89.8±4.3 | 0.00181 |
|  |  |  | 0.04 | 254 | - | - |
|  |  |  | 0.01 | 64 | - | - |

A quantitative assessment of the effects of the antagonist hits on AgamORco channel function was undertaken through measurements of reductions in the OA-dependent channel activity due to the presence of increasing quantities of the antagonists relative to the previously reported values for IPC (34). As is depicted by the dose response curves presented in **Figure 3**, all 5 hits were found to antagonize AgamORco channel function in a dose-dependent manner with IC_50_ values ranging from 23 to 83 μM.

**Figure 3.**
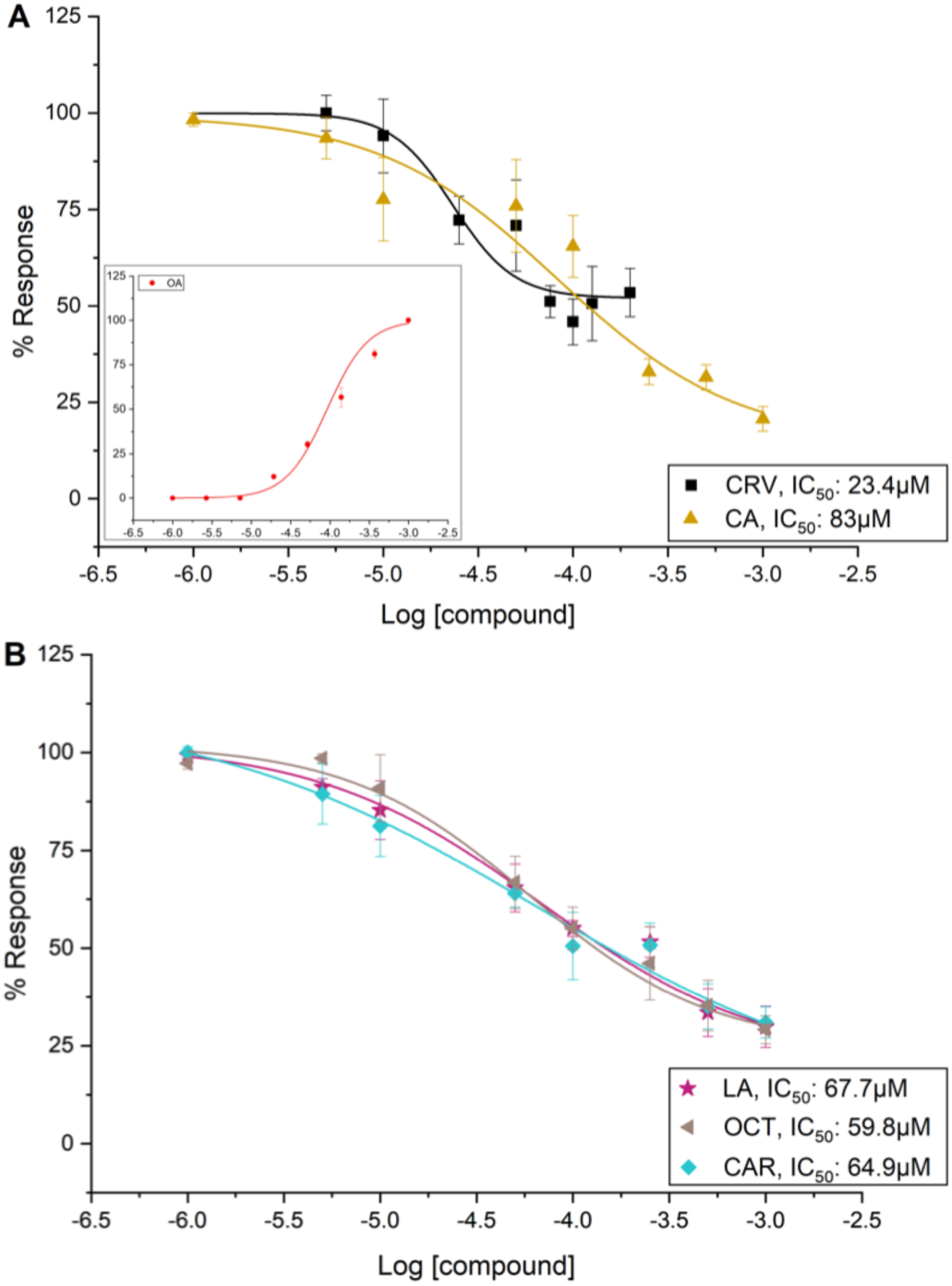
Dose-dependent inhibition of AgamORco function by identified antagonists. IC_50_ values for all tested compounds except CRV were determined using antagonist concentrations in the range of 1μM to 1mM. Because CRV was found to be toxic to the cells at concentrations above 250 μM, concentrations ranging from 1 to 200 μM were used for its dose-response evaluation. ***Panel A:*** The IC_50_ values determined for the two known repellents, CRV and CA, were 23.4 μM (pIC_50_: 4.63179±0.09524, R^2^: 0.9755) and 83 μM (pIC_50_: 4.08243±0.28481, R^2^: 0.99989), respectively, while that for the previously characterized ORco antagonist IPC was 41.7 μM (pIC_50_: 4.37919±0.061, R^2^: 0.9883, Tsitoura et al., 2015). The EC_50_ for the OA ORcoRAM2 from the curve that is shown in the inset is 91.9 μM (pIC_50_: 4.03684±0.11567, R^2^: 0.99998). ***Panel B:*** The IC_50_ values for the 3 new putative ORco antagonists, LA, OCT and CAR, were 67.7 μM (pIC_50_: 4.16927 ±0.15954, R^2^: 0.99999), 59.8 μM (pIC_50_: 4.22309, R^2^: 0.99988) and 64.9 μM (pIC_50_: 4.18725 ±0.31571, R^2^: 0.99998), respectively. Error bars indicate mean ± SE. Data points were normalized to the maximum value and multiplied by 100.

To confirm the cross-species bioactivity of these compounds in mosquitoes, we tested the activity of the two most potent antagonists, CRV and OCT (**Figure 3**) on *Xenopus* oocytes expressing *Ae. aegypti* ORco (AaegORco). While neither of the two compounds elicited currents in water-injected oocyte controls (**Figure S-1**), CRV and OCT reduced VUAA1-activated currents (**Figure 4A**) by approximately 60% and 85%, respectively (**Figure 4B**), in accordance with the insect cell-based results where CRV was found to be a more potent inhibitor compared to OCT (**Figure 3**).

**Figure 4.**
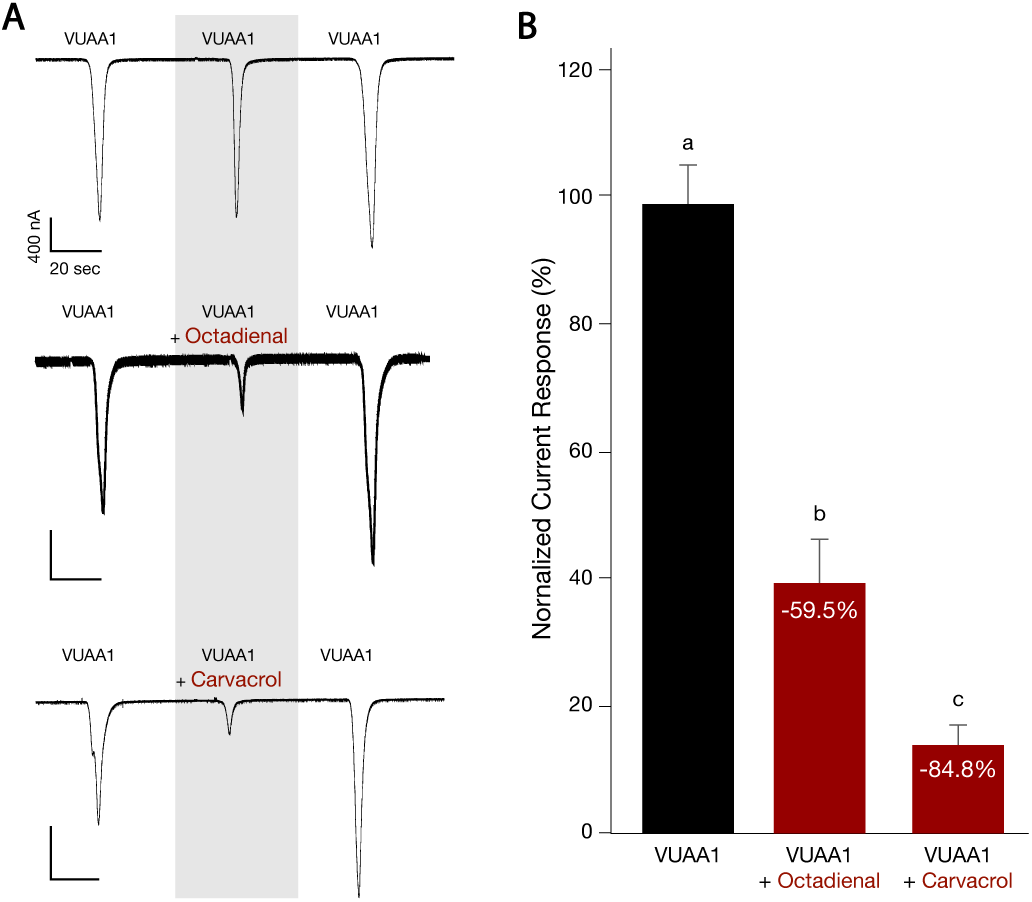
Octadienal and carvacrol inhibit VUAA1-induced ORco function in frog oocytes. ***Panel A*:** Representative current traces of oocytes expressing AaegORco following exposure to 200 μM VUAA1 alone or in combination with equimolar concentrations of octadienal or carvacrol. ***Panel B*:** Normalized responses of AaegORco to VUAA1 alone or in combination with octadienal or carvacrol. Odorants effects were statistically significant (one-way ANOVA followed by Tukey’s post-test; P<0.0001; mean responses ± s.e.m.; VUAA1 alone, n=7; OCT and CRV, n=6).

### Natural VOCs acting as AgamORco antagonists repel *Aedes albopictus* mosquitoes

The five AgamORco antagonists identified in the cell-based screening assay were tested subsequently for repellence activity against laboratory populations of *Ae. albopictus* (also known as the “Asian tiger mosquito”), an aggressive mosquito species, based on the reduction in the numbers of landings on an exposed portion of a human hand. The widely used repellent DEET (45) and the strong mosquito repellent IPC (44,46), previously characterized as AgamORco antagonist (34), were used as standards.

The bioassays (see **Table S-2** for quantifications of landing numbers) showed that at the highest tested dose (0.2 μL/cm^2^, 1-1.4 μmole/cm^2^), all compounds significantly reduced mosquito landing counts relative to the solvent controls (**Figure 5A** and **Table 1**). In the presence of 0.04 μL/cm^2^ (210-280 nmole/cm^2^), the strongest ORco antagonists, CRV and OCT, displayed repellent activities comparable to DEET, while the activity of CA was noticeably lower (**Figure 5B** and **Table 1**). At the lowest tested dose of (0.01 μL/cm^2^, 52-70 nmole/cm^2^), all compounds were found to display repellent activities lower than DEET, with OCT eliciting the lowest repellency of all (**Figure 5C** and **Table 1**).

**Figure 5.**
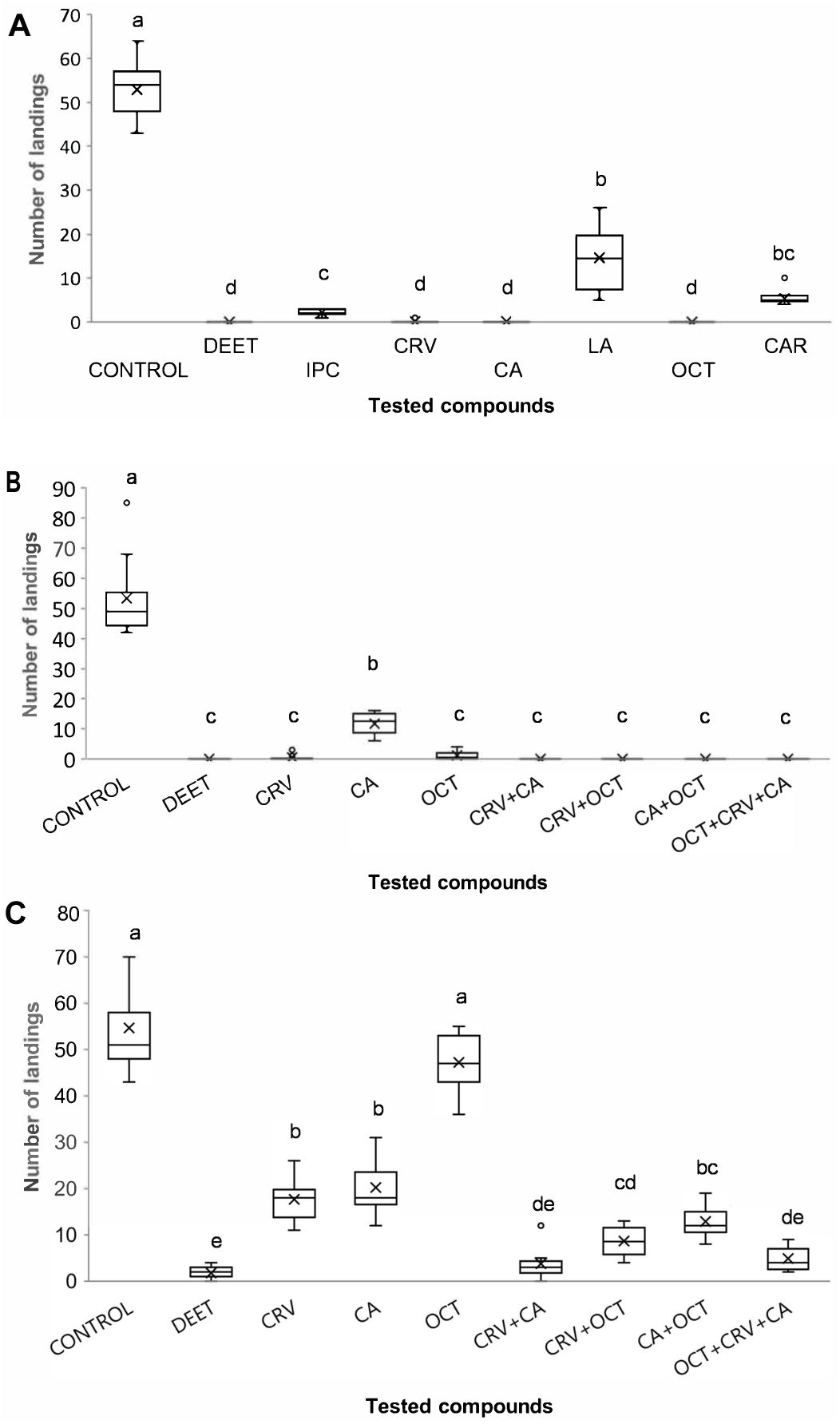
Repellent activities of ORco antagonists. Box plots depicting landings distributions of *Ae. albopictus* mosquitoes after exposure to a dose of 0.2 (**panel A**), 0.04 (**panel B**) or 0.01 (**panel C**) μL/cm^2^ (for molar concentrations see also **Tables 1** and **2**) of tested material for five minutes. On the right half of panels B and C, the effects of binary and ternary mixtures are presented. Different letters, a-e, indicate significant differences among treatments (P<0.05, Mann–Whitney U test with Bonferroni correction).

### Binary mixtures of ORco antagonists are more active than single compounds

The three compounds with the strongest repelling indices were subsequently tested in the bioassay as binary and ternary mixtures, using two different doses, medium and low (**Figure 5B** and **5C**, respectively, and **Table 2**).

**Table 2:** Low doses of compound mixtures are more active repellents than single ones. All tests were carried out over a period of 5 minutes. Dichloromethane (DCM) was used as a control and the number of landings for the experiments involving total compound doses equivalent of 0.01 μL/cm^2^ are presented in **Supplementary Table 2**. Statistically significant differences are those with Pr(<[0.05]).

| Compound | $\mu\text{L}$ equivalent of each compound per $\text{cm}^2$ | nmole of each compound per $\text{cm}^2$ | mean % repellency | DEET $\text{Pr}(<[t])$ | Single compound $\text{Pr}(<[t])$ | | |
| --- | --- | --- | --- | --- | --- | --- | --- |
| DEET | 0.01 | 52 | 96.7±2.6 | | | | |
| CRV | 0.01 | 65 | 67.8±9.0 | 6.9E-16 | | | |
| CA | 0.01 | 65 | 63.2±11.8 | 1.3E-14 | | | |
| OCT | 0.01 | 70 | 13.8±12.8 | 8.6E-25 | | | |
| CRV+CA | 0.005 | 32.5+32.5 | 93.1±6.8 | 0.02971 | **CRV** | **CA** | |
|  |  |  |  |  | 1.8E-05 | 3.7E-05 |  |
| CRV+OCT | 0.005 | 32.5+35 | 84.2±6.6 | 1.3E-09 | **CRV** | **OCT** | |
|  |  |  |  |  | 0.00087 | 3.9E-09 |  |
| CA+OCT | 0.005 | 32.5+35 | 76.5±6.9 | 8.0E-14 | **CA** | **OCT** | |
|  |  |  |  |  | 0.02423 | 8.3E-08 |  |
| CRV+CA+OCT | 0.0033 | 21.7+21.7+23.3 | 91.1±5.2 | 0.00019 | **CRV** | **CA** | **OCT** |
|  |  |  |  |  | 4.3E-05 | 9.6E-05 | 4.5E-09 |

At the lowest antagonist doses examined (0.01 μL/cm^2^), equivolume binary mixtures consisting of 0.005 μL/cm^2^ each of CRV (32.5 nmole/cm^2^) and CA or OCT (32.5 and 35 nmole/cm^2^, respectively), reduced mosquito landing rates by 93.1% (CRV+CA), 84.2% (CRV+OCT) and 76.5% (CA+OCT), respectively, which were significantly higher than landing rates elicited by single compounds administered at 0.01 μL/cm^2^ (67.8%, 63.2% and 13.8% for CRV, CA and OCT, respectively; **Figure 5C** and **Table 2**).

The CA-CRV mixture, in particular, which displayed repellent activity against *Ae. albopictus* very similar to DEET (landing inhibition of 93.1% versus 96.7%), was previously shown to repel *An. gambiae* and *Culex spp* mosquitoes in the field more efficiently than DEET (33). A ternary mixture consisting of 0.0033 μL/cm^2^ each of CRV (21.7 nmole/cm^2^), CA (21.7 nmole/cm^2^) and OCT (23.3 nmole/cm^2^), displayed repellence activity of 91.1% against *Ae. albopictus*, comparable to that of the CRV+CA binary mixture at the same total antagonist amount of 65 nmole/cm^2^ (93.1%).

### Binding competition assays reveal possible mechanism for enhanced activity of binary mixtures

A possible explanation for the increased repellent activity of binary antagonist mixtures is that it may be caused by their combined interactions within the ORcoRAM2 binding site (47) or with additional, distinct binding sites. Such interactions could impose enhanced conformational rearrangements in ORco, steric hindrance in the agonist binding site and enhanced inhibition of ORco function. Moreover, given the small size of at least some of the identified antagonists, it is also possible that a single binding site, e.g., the one to which the ORco agonist binds, could accommodate the binding of two different antagonists that act in an additive fashion thus causing a higher degree of ORco inhibition relative to the single ones.

To distinguish between competitive (orthosteric) and non-competitive (allosteric) effects, we carried out antagonist binding competition experiments. These assays involved antagonist and agonist binding to AgamORco that could distinguish between competitive and non-competitive binding of the examined antagonists relative to the agonist binding site. The results of these competition experiments are shown in **Figure 6**.

**Figure 6.**
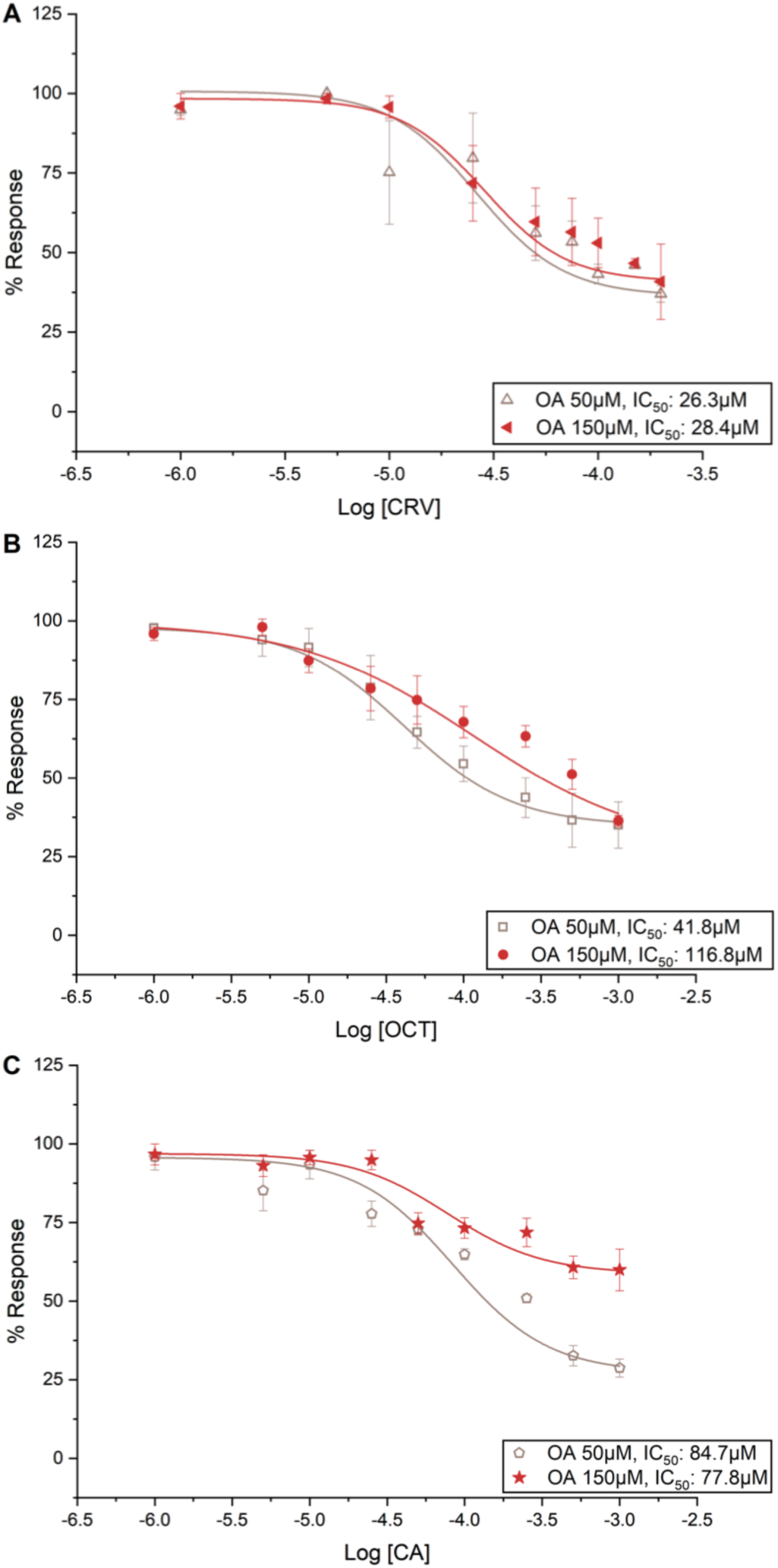
ORco competition assays. ***Panel A:*** The IC_50_ values determined for carvacrol (CRV) in the presence of 50 μM and 150 μM ORcoRAM2 were 26.3 μM (pIC_50_: 4.58001±0.21732, R^2^: 0.99998) and 28.4 μM (pIC_50_: 4.54608, R^2^: 0.99747), respectively. There is no noticeable shift in IC_50_ concentration as OA concentration increases (IC_50_ of CRV with 100 μM OA is 23.4 μM), expected when compounds bind at different binding sites, with allosteric antagonistic effect. ***Panel B:*** The IC_50_ values determined for cumin alcohol (CA) in the presence of 50 μM and 150 μM ORcoRAM2 were 84.7μM (pIC_50_: 4.07226±0.23144, R^2^: 0.99998) and 77.8μM (pIC_50_: 4.10898±0.17809, R^2^: 0.99984), respectively. ***Panel C:*** Similarly, the IC_50_ values determined for octadienal (OCT) in the presence of 50 μM and 150 μM ORcoRAM2 were 41.8μM (pIC_50_: 4.37887±0.03444, R^2^: 0.99999) and 116.8μM (pIC_50_: 3.93263±0.26793, R^2^: 0.99973), respectively. There is a dextral shift of the curve and a concomitant increase of the IC_50_ as the OA concentration is increased (=3 fold), expected when both compounds compete for the same binding site. Error bars indicate mean ± SE. Data points were normalized to the maximum value.

These experiments suggest that CRV, with very similar IC_50_ values, 26.3 and 28.4 μM in the presence of 50 and 150 μM Orco agonist, respectively (**Figure 6A**), is a non-competitive (allosteric) antagonist of the ORco agonist. In contrast, with decreasing potency in the presence of increasing OA amounts (IC_50_ of 41.8 and 116.8 μM in the presence of 50 and 150 μM of ORcoRam2, respectively; **Figure 6B**), OCT appears to be a competitive (orthosteric) inhibitor of ORcoRAM2. Cumin alcohol, on the other hand, also behaves as an allosteric inhibitor of ORco function as its IC_50_ values against 50 and 150 μM ORcoRAM2 is maintained at similar levels, 84.7 and 77.8 μM, respectively (**Figure 6C**). However, relative to CRV, CA is a less potent antagonist, as its ability to antagonize the effect of 150 μM ORcoRAM2 is reduced.

## DISCUSSION

OBPs and ORs expressed predominantly in female mosquitoes are known to constitute promising targets for the discovery of molecules capable of altering the odor sensing capacity and odor-evoked behaviors of mosquitoes (48). Previous work has shown that some strong mosquito repellents of natural origin (33) act as ORco antagonists (34). Moreover, ORco-specific synthetic agonists, such as VUAA1 and OrcoRAM2, were shown to activate ORx/ORco channels in cultured insect cells (34) and to also act as positive allosteric modulators (PAMs) of odorant receptor function (35). Thus, ORco is a rational target for molecules that could function as modulators of peripheral olfactory functions in mosquitoes and, probably, other insect species as well. Consequently, the employment of screening platforms that exploit the capacity of ORco to form functional homomeric ion channels in cultured cells and the identification of specific ORco agonists or antagonists, may result in the discovery of natural or synthetic modulators of olfaction-dependent mosquito behaviors. Such modulators may be either “anosmia”-inducing factors or olfactory enhancers.

In this report, we have presented an integrated study that included the design and use of a convenient screening platform that allows detection of the presence of ORco functional modulators in collections of metabolites of natural or synthetic origin as well as pharmacological and functional aspects of identified modulators. Specifically, the screening of the 50 natural metabolites whose common property has been their volatility, resulted in the detection of three novel AgamORco antagonists, LA, OCT and CAR (**Figure 2 and 3**). Two additional compounds, CRV and CA, which were previously shown to inhibit the function of AgamORco and several AgamORx/ORco receptors (34), were confirmed here to be AgamORco antagonists and further characterized pharmacologically (**Figures 2-4**). Interestingly, the identified antagonists are not characterized by the presence of a single functional group. Thus, CRV and CA are aromatic ligands decorated with electrophilic functionalities, while LA and OCT are similarly decorated aliphatic compounds. On the other hand, CAR is a bicyclic non-polar molecule. As discussed below, such structural features may relate to the nature of the binding sites of these compounds on ORco.

Our earlier findings concerning the bioactivities of CRV and CA, now confirmed to be AgamORco antagonists, which were shown to repel effectively *An. gambiae* and *Culex spp*. mosquitoes (33), raised the question of whether these as well as the new antagonists identified in this study were also active against *Aedes*, the third major mosquito genus of the *Culicidae* family, which comprises multiple haematopagous species and infectious disease vectors. The initial testing of two selected antagonists, CRV and OCT, in *X. laevis* oocytes expressing ORco of *Ae. aegypti* revealed significant inhibition of AaegORco function (**Figure 4**). In addition, all identified antagonists were shown to repel *Ae. albopictus* mosquitoes to various degrees (**Table 1** and **Figure 5**). The combined results constitute proof of principle for the notion that AgamORco antagonists are efficient blocking agents of olfactory function in multiple mosquito genera and that the search for new compounds capable of interfering with mosquito olfactory functions by screening VOC collections for ORco, as opposed to multiple ORx-specific antagonist activities, is feasible.

A recent study involving the functional screening of a small collection of commercially available natural compounds selected through machine learning methodologies in *X. laevis* oocytes expressing AgamOrco, identified two AgamOrco antagonists, which inhibited odorant responses in electroantennogram and single sensillum recordings of adult *Drosophila melanogaster* antennae and inhibited odorant-directed behaviors in larvae of the same species (49). Interestingly, this study, whose results are concordant with ours with respect to the cross genus bioactivities of AgamORco antagonists, identified linalyl formate, a compound with a structure almost identical to that of LA, as one of the two AgamORco antagonists that inhibited odorant-directed behaviors in *Drosophila* larvae.

The combined findings on the physiological and behavioral effects of AgamORco antagonists on different dipteran species are obviously due to the very high phylogenetic conservation of ORco amongst insect species (10-14). In this regard, we note that the notion that agonists capable of activating constitutively the common subunit of odorant receptors should cause olfactory confusion has also been proposed (25). However, relevant behavioral experimentation to confirm this notion have yet to be presented.

An obvious aspect that needs to be explored further concerns the molecular mechanism underlying the behavioral effects of the identified AgamORco antagonists on the targeted mosquitoes. Although explanations involving ORco-independent pathways may be invoked to explain the behavioral changes induced in mosquitoes exposed to these volatile antagonists, our results are consistent with the hypothesis that the observed effects are due to the functional inhibition of the olfactory apparatus caused by their direct binding to the obligatory ORco subunit of odorant receptors. Thus, the inhibitors of ORco homomeric channels formed in cultured cells apparently become common intraspecific inhibitors of essentially all ORx/ORco heteromeric receptors *in vivo*, in a way analogous to but much broader than the recently proposed intraspecific inhibitors of heteromeric receptors (50). Based on the proposed ability of the identified ORco-targeting VOCs to inhibit the function of multiple ORs, we consider it likely that they cause anosmia-like effects to the targeted mosquitoes. Proof of this hypothesis will have to await the identification of the specific antagonist binding sites and detailed mutagenesis studies. The proposed mode of action for the identified bioactive VOCs is thus distinct from the receptor-independent function of DEET, which, apparently, “repels” mosquitoes and other insects, at least in part, via its association with volatile receptor ligands acting as attractants, thereby reducing their volatility and effective concentrations thus masking their presence in the mosquito’s environment (51).

An additional noticeable finding of this study has been the enhanced repellent action of binary combinations of ORco antagonists on the behavior of *Ae. albopictus* adults. Such effects had been noted previously in our studies on laboratory and field populations of *An. gambiae* and *Culex spp* (33). The enhanced bioactivities of binary mixtures of chemically diverse AgamORco antagonists raised the question of the possible relevance of antagonist binding sites. The binding competition assays for CRV and CA against ORcoRAM2 (**Figure 6A** and **6B)** suggested that the two compounds, which maintain very similar IC_50_ values in the presence of 50 or 150 μM (as well as 100 μM; **Figure 3A**) of OA, are binding to sites different than the one to which ORcoRAM2 binds and are, therefore, non-competitive, allosteric antagonists of the proposed OA binding sites on the AgamORco tetramer (47). The enhanced performance of the CRV+CA mixture in the mosquito landing inhibition assays relative to CRV or CA alone (**Table 2**), further suggests separate CRV and CA binding sites despite the apparent structural similarities between these two compounds. The alternative possibility that these two compounds synergize by binding to different sections of a common allosteric binding pocket, cannot be excluded without further experimentation. In contrast, the binding competition assays for OCT (**Figure 6C**) suggest that this compound is a competitive, orthosteric antagonist of AgamORco with respect to its agonist (ORcoRAM2) binding site. Accordingly, we are attributing the synergistic effects of the CRV+OCT and CA+OCT mixtures relative to CRV, CA or OCT alone to the simultaneous binding of an orthosteric and an allosteric antagonist on AalbORco. Although the antagonist binding competition experiments provide clues related to the nature of their binding sites relative to the postulated one for ORcoRAM2 (47), the precise nature of the binding sites, particularly for CRV and CA remain to be determined. Molecular Dynamics and molecular docking studies are currently in progress in an effort to identify candidate binding sites in the recently resolved homotetrameric complex of the ORco channel (47).

The cross-species bioactivities of compounds with mosquito repelling activities discovered through the AgamORco VOC screen, which, due to the high conservation of ORco across phylogeny, are also capable of repelling, thus offering biting protection from other insects and arachnoids such as *Lutzomyia longipalpis* sandflies and *Ixodes ricinus* ticks (52), may raise concerns regarding their environmental safety. In this respect, it should be stressed that such repellent compounds pose no danger to the environment, as they are destined to be used on a limited scale, either for personal protection or as spatial repellents.

## EXPERIMENTAL PROCEDURES

### Mosquitoes

Adult *Aedes albopictus* mosquitoes were obtained from a laboratory colony maintained at 25±2 °C, 80% relative humidity, and 16/8-h light/dark photoperiod at the Benaki Phytopathological Institute, Kifissia, Greece (53). Plastic beakers with 100 ml water and strips of moistened filter paper were inserted in the cages for oviposition. The eggs were kept damp for a few days and then placed in enamel pans for hatching. The larvae were reared in tap water-filled cylindrical enamel pans, approximately 400 larvae per pan, and were fed *ad libitum* with powdered fish food (JBL Novo Tom 10 % Artemia) until the emergence of adults. Adult mosquitoes were collected periodically with a mouth aspirator and transferred to the rearing cage. Females were fed with fresh chicken blood using a Hemotek^©^ blood feeding system (54).

### Chemicals

The ORco agonists, repellents and the 50 VOCs analyzed in the current study are presented in **Table S-1**. Carvacrol (hereafter CRV), linalyl acetate (hereafter LA), (2*E*,4*E*)-2,4-octadienal (hereafter OCT) and (1S)-3-carene (hereafter CAR) were purchased from Sigma Aldrich, isopropyl cinnamate (hereafter IPC) from Alfa Aesar, cumin alcohol (hereafter CA) from Acros Organics, *N*-(4-ethylphenyl)-2-{[4-ethyl-5- (3-pyridinyl)-4H-1,2,4-triazol-3-yl]thio}acetamide (ORco Receptor Agonist Molecule 2, hereafter ORcoRAM2) from Asinex Corporation and Vitas M Chemical Ltd, *N*-(4-ethylphenyl)-2-{[4-ethyl-5-(3-pyridinyl)-4H-1,2,4-triazol-3-yl]thio}acetamide (hereafter VUAA1) from Innovapharm Ltd., *N,N-*diethyl-3-methylbenzamide (hereafter DEET) from Sigma-Aldrich and coelenterazine from Biosynth. Initial stock solutions (50 mM) for ORcoRAM2 and VUAA1 were prepared in DMSO and stored at - 20°C, while initial stocks of VOCs and coelenterazine were prepared in ethanol as needed and stored at -20°C. For the insect cell-based screening assay, working solutions were prepared in modified Ringer’s buffer (25 mM NaCl, 190 mM KCl, 3 mM CaCl_2_, 3 mM MgCl_2_, 20 mM Hepes, 22.5 mM glucose, pH 6.5; 35), so that final DMSO concentrations did not exceed the range of 0.2-0.35%.

### Transformation of Bm5 cells for AgamORco and Photina expression and Ca^***2+***^ ***influx assays***

The screening platform shown in **Figure 1** was employed as tool for discovery of new compounds capable of modulating mosquito ORco activity and olfaction-mediated behaviors. It consists of lepidopteran cultured cells (*Bombyx mori* Bm5; 55) expressing constitutively AgamORco, which forms a ligand-gated ion channel (34,35), and Photina™ (56), a reporter photoprotein activated by Ca^2+^ ions entering the cells upon activation of the ORco channel. Briefly, Bm5 cells were stably transformed to express the cDNAs for AgamORco and the reporter photoprotein Photina™ from high expression level pEIA plasmid vectors (57-60) as previously described (21,34). Cell lines were maintained at 28°C and grown in IPL-41 insect cell culture medium (Genaxxon Bioscience GmbH) supplemented with 10% fetal bovine serum (Biosera) in the presence of 10 μg/mL puromycin. Ligand binding to the ORco channel and subsequent functional effects were monitored using Photina™ as Ca^2+^ influx biosensor. Briefly, cells were resuspended in modified Ringer’s buffer, seeded in a 96-well plate (200,000-300,000 cells/well) and incubated with 5 μM coelenterazine for two hours at room temperature in the dark. Baseline and maximum luminescence outputs, obtained by addition of 1% Triton-X100, were recorded in an Infinite M200 microplate reader (Tecan) at 4 s intervals for up to 20 s. The cells were subjected to two cycles of compound additions. Initially, a tested compound was added at 100 μM and the ORco channel response was monitored. Cells were allowed to return to baseline luminescent and this was followed by addition of 100 μM of the ion channel-activating OA and measuring the secondary effect of ligand binding in terms of luminescence emission (4 s intervals for 80 s). Initial luminescence data were acquired using i-Control 1.3 software by Tecan. Relative luminescence values were normalized by considering ORco agonist luminescent response as the maximal (100%) receptor response for each set of experiments. Each independent experiment was run in triplicate and repeated at least three times.

### Binding competition assays

Cultured lepidopteran cells expressing AgamORco and Photina™ were resuspended in modified Ringer’s medium, seeded in 96-well plates and incubated with 5μM coelenterazine for two hours at room temperature in the dark. A previously tested compound was then added at concentrations ranging from 1 μM to 1 mM and the induced luminescence was measured. This was followed by addition of ORcoRAM2 in all wells at final concentrations of 50 or 150 μM. The dose-dependent effects of ORco activation inhibition was monitored via luminescence reduction, relative to the controls. Each independent experiment was run in triplicate and repeated at least three times. Curve fitting and EC_50_/IC_50_ value calculations were performed using OriginPro 8 software by OriginLab Corporation.Dose response curves were plotted by fitting the normalized data into the equation, 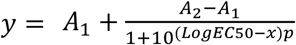 where A_1_ and A_2_ are the bottom and top asymptotes respectively, *p* is the Hillslope, *y* is the % response at a given concentration and *x* is logarithm of ligand concentration.

### Expression of AaegORco in Xenopus laevis oocytes and electrophysiological recordings

*In vitro* transcription and two-microelectrode voltage-clamp electrophysiological recordings were performed as described previously (61). Briefly, ORco of *Aedes aegypti* (AaegORco) was synthesized using the mMESSAGE mMACHINE® SP6 kit (ThermoFisher Scientific) from the linearized pSP64tRFA expression vector. Harvested *X. laevis* oocytes were manually separated from the ovaries prior to collagenase treatment (8mg/mL, 30min, 18°C) in order to remove the follicular layer. Stage V-VI oocytes were rinsed in washing solution (96 mM NaCl, 2mM KCl, 5mM MgCl_2_ and 5mM HEPES, pH 7.6) and microinjected with 1 μL of AaegOrco (3 μg/μL) and 2 μL of double-distilled water. Injected oocytes were incubated at 18°C for 3 days in Ringer’s solution (96 mM NaCl, 2 mM KCl, 5 mM MgCl_2_, 0.8 mM CaCl_2_ and 5 mM HEPES, pH 7.6) supplemented with 5% dialyzed horse serum, 50 μg/mL tetracycline, 100 μg/mL streptomycin and 550 μg/mL sodium pyruvate. Whole-cell currents were recorded using the two-microelectrode voltage-clamp technique. During recording sessions, the holding potential was maintained at - 80 mV using an OC-725C oocyte clamp (Warner Instruments, LLC, Hamden, CT, USA). Oocytes were placed in a RC-3Z oocyte recording chamber (Warner Instruments, LLC, Hamden, CT, USA) and exposed for 8 seconds to 200 μM VUAA1 (Innovapharm Ltd., Kiev, Ukraine), 2,4-Octadienal predominantly trans (Sigma-Aldrich) or carvacrol (Sigma-Aldrich). Currents were allowed to return to baseline between odorant applications. Data acquisition and concentration-response analyses were carried out with a Digidata 1550A and the pCLAMP10 software (Molecular Devices, Sunnyvale, CA, USA). Statistical significance was evaluated with one-way ANOVA followed by Tukey’s post-test.

### Repellence Bioassays

For the *in vivo* determination of the repellent activity of VOCs, the assessment was based on human hand landing counts (62). The study was conducted using cages (33×33×33 cm) equipped with a 32×32 mesh at one side, each containing 100 5- to 10-day old adult mosquitoes (sex ratio, 1:1) starved for 12 h at 25±2 °C and 70–80% relative humidity. A volunteer’s hand covered by a plastic glove with a dorsal side opening measuring 5×5 cm was employed for all bioassays. Tested compounds were applied on chromatography paper (Whatman), over a 24 cm^2^ total area, at three doses equivalent to 0.2, 0.04 and 0.01 μL/cm^2^ (1000-1400, 200-280 and 50-70 nmole/cm^2^, respectively, depending on compound molecular mass; see also **Table 1**) diluted with dichloromethane (DCM). Control experiments with compound-free DCM solvent or DEET treatments (negative and positive controls, respectively) were included as standards. Each treatment was repeated 8 times on four human volunteers. Replicate experiments were n=15 for the solvent- and DEET-treated controls. The effects of tested ORco antagonists on *Ae. albopictus* landings was estimated using the Kruskal–Wallis test (63). When significant differences were detected, Mann–Whitney U tests with Bonferroni correction (64) were carried out for pair-wise among treatments. Mosquito landings for each treatment were counted over 5-minute periods. Landing numbers were converted to repellence indices (RI±SE) using the equation 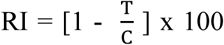, where *C* was the number of landings in control and *T* the number of landings in treatment.

## ACKNOWLEDGMENTS

We thank C. Meristoudis for technical assistance with the generation of stable cell lines at the initial stages of the project.

## Author Contributions

KI, AM, JDB contributed conception, design and supervision of the study; SS organized the compound database and provided compounds; GK, NS, PT, VK, DMR, EY performed investigation experiments and interpreted results; DPP provided methodology, data analysis and curation; GK and DPP performed statistical analyses; GK and KI wrote the first draft of the manuscript; PT, AM, JDB and SS wrote sections of the manuscript. All authors contributed to manuscript review and editing, read and approved the submitted version.

